# Poly-Exposure and Poly-Genomic Scores Implicate Prominent Roles of Non-Genetic and Demographic Factors in Four Common Diseases in the UK

**DOI:** 10.1101/833632

**Authors:** Yixuan He, Chirag M Lakhani, Arjun K Manrai, Chirag J Patel

## Abstract

While polygenic risk scores (PRSs) have been shown to identify a small number of individuals with increased clinical risk for several common diseases, non-genetic factors that change during a lifetime, such as lifestyle, employment, diet, and pollution, have a larger role in clinical prediction. We analyzed data from 459,613 participants of the UK Biobank to investigate the independent and combined roles of demographics (e.g., sex and age), 96 environmental exposures, and common genetic variants in atrial fibrillation, coronary artery disease, inflammatory bowel disease, and type 2 diabetes. We develop an additive modelling approach to estimate and validate a poly-exposure score (PXS) that goes beyond consideration of a handful of factors such as smoking and pollution. PXS is able to identify groups with high prevalence of the four common disease comparable to, if not better, than the PRS. Type 2 diabetes has the largest discrepancy in PXS and PRS performance, defined as the maximum area under the receiver-operator curve (AUC) (PXS AUC of 0.828 [0.821-0.836], PRS AUC of 0.711 [0.702-0.720]). Most importantly, we show that PXS identifies individuals that have low genetic risk but high overall risk for disease. While PRS is useful for screening genetically exceptional individuals in a time-invariant way, broader consideration of multiple non-genetic and modifiable factors is required to fully translate risk scores to the bedside for precision medicine. All results and the PXS calculator can be found in our web application http://apps.chiragjpgroup.org/pxs/.

## INTRODUCTION

Recent advances in sequencing technologies have led to an explosion of genome-wide association studies (GWASs) that have identified thousands of genetic loci associated with complex traits and diseases in humans^1^. At the same time, it has become widely accepted that human traits and diseases are also heavily influenced by environmental or non-genetic factors. However, studies of non-genetic exposure factors often only consider a single or handful of factors at a time^1^. Further still, most research is siloed, examining either genetics, demographics (e.g., age and sex), or exposure variables associated with disease. The relative predictive power of basic demographic and non-genetic variables alone (and together) is mostly unknown. With a few exceptions^2^, rarely have investigations considered multiple non-genetic and genetic factors simultaneously, possibly due to the lack of data resources which have both genetic information and a large set of environmental exposure variables.

While genome-wide polygenic risk scores (PRS) have been shown to identify individuals with significantly increased clinical risk for several common diseases^3,4^, a comparable study on combined multiple exposure/non-genetic and demographic factors does not exist. For example, Park et al. report an “environmental risk score” for serum lipid (total cholesterol, high-density lipoprotein cholesterol (HDL), low-density lipoprotein cholesterol (LDL) and triglycerides), but only pollutants are considered in the score^5^. On the other hand, “lifestyle” scores to measure modifiable factors can be used (e.g. Khera et al.^6^), but it is a challenge to analytically define “lifestyle”^7^. Furthermore, it is unclear if the combined poly-exposure additive effect can outperform any single exposure alone or genetic variants for predicting risk of common diseases.

Thus, we term the “poly-exposure score” (PXS) and the “poly-demographic score” (PDS). Using self-reported and hospital admission information from 459,613 total participants of the UK Biobank, we developed PXSs and PDSs that combines up to 96 independent exposure and 42 demographic variables for atrial fibrillation (AF), coronary artery disease (CAD), inflammatory bowel disease (IBD), and type 2 diabetes (T2D). We demonstrate the ability of both PXS and PDS to identify groups with high prevalence of disease and highlight the importance to consider both genetic and non-genetic factors in genetics research and potentially clinical care.

## RESULTS

### Poly-exposure score captures more information than any environmental exposure alone

In summary, we conducted an eXposure-Wide Association Study (XWAS) with 96 exposure variables in the training set of 104,623 individuals (Figure 1, Methods)^7–9^. These exposure variables include indicators of alcohol, air pollution, noise pollution, dietary factors, dietary nutrients, early life factors, education levels, employment status, household information, physical activity, sleeping habits, smoking, and area-level indicators of a geography/tract, including Townsend index (See Supplementary Table 1 for the entire list). We refer to the entire array of potential environmental or non-genetic factors as ‘exposures’.

**Figure 1:**
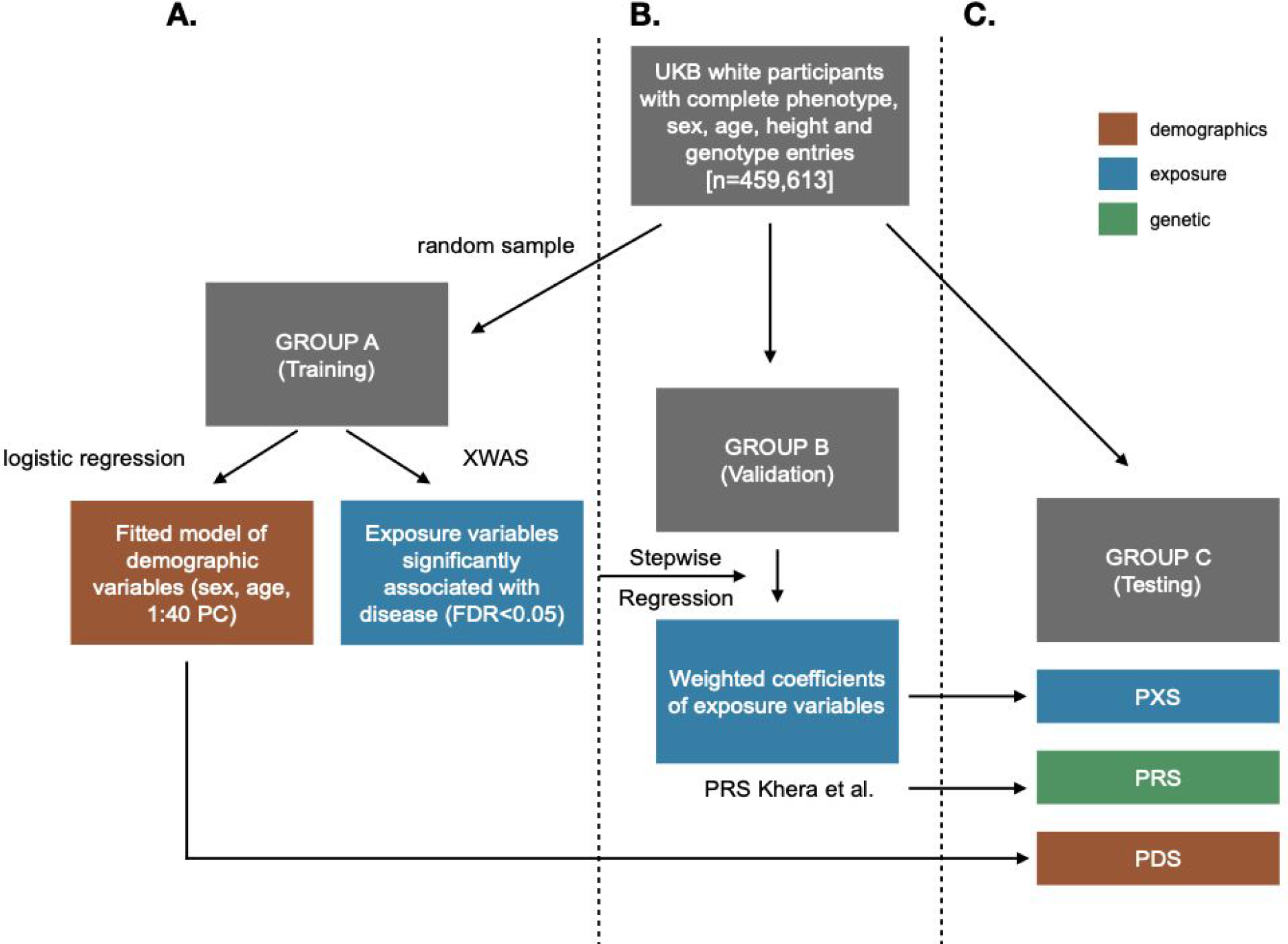
Study design. White participants with complete demographic data (n=459,613) were randomly divided into training, validation, and testing sets (cohorts A, B and C). A) For each of the four diseases, the initial univariate XWAS analysis and fitted demographic model were conducted in cohort A. B) Exposure variable selection was conducted in the cohort B. C) Testing the performance of all three scores was conducted in cohort C. (PDS: poly-demographic score, PRS: polygenic risk score, PXS: poly-exposure score.)

Depending on the amount of missing information, each XWAS regression analysis had a different sample size, with a mean of 102,911 individuals (*se*=232 individuals) (Supplementary Table 1). There were 37 significantly associated exposure variables with AF, 49 with CAD, 19 with IBD, and 74 with T2D (Figure 2A). In AF, CAD, and T2D, “Major dietary changes in the last 5 years: Yes, because of illness”, “Usual walking pace: Slow”, and “Time spend watching TV” were among the most significantly associated responses and were positively associated with the diseases. In IBD, “Major dietary changes in the last 5 years: Yes, because of illness”, “Never eat dairy products”, “Getting up in the morning: not very easy” were the top most significant positively associated responses.

**Figure 2:**
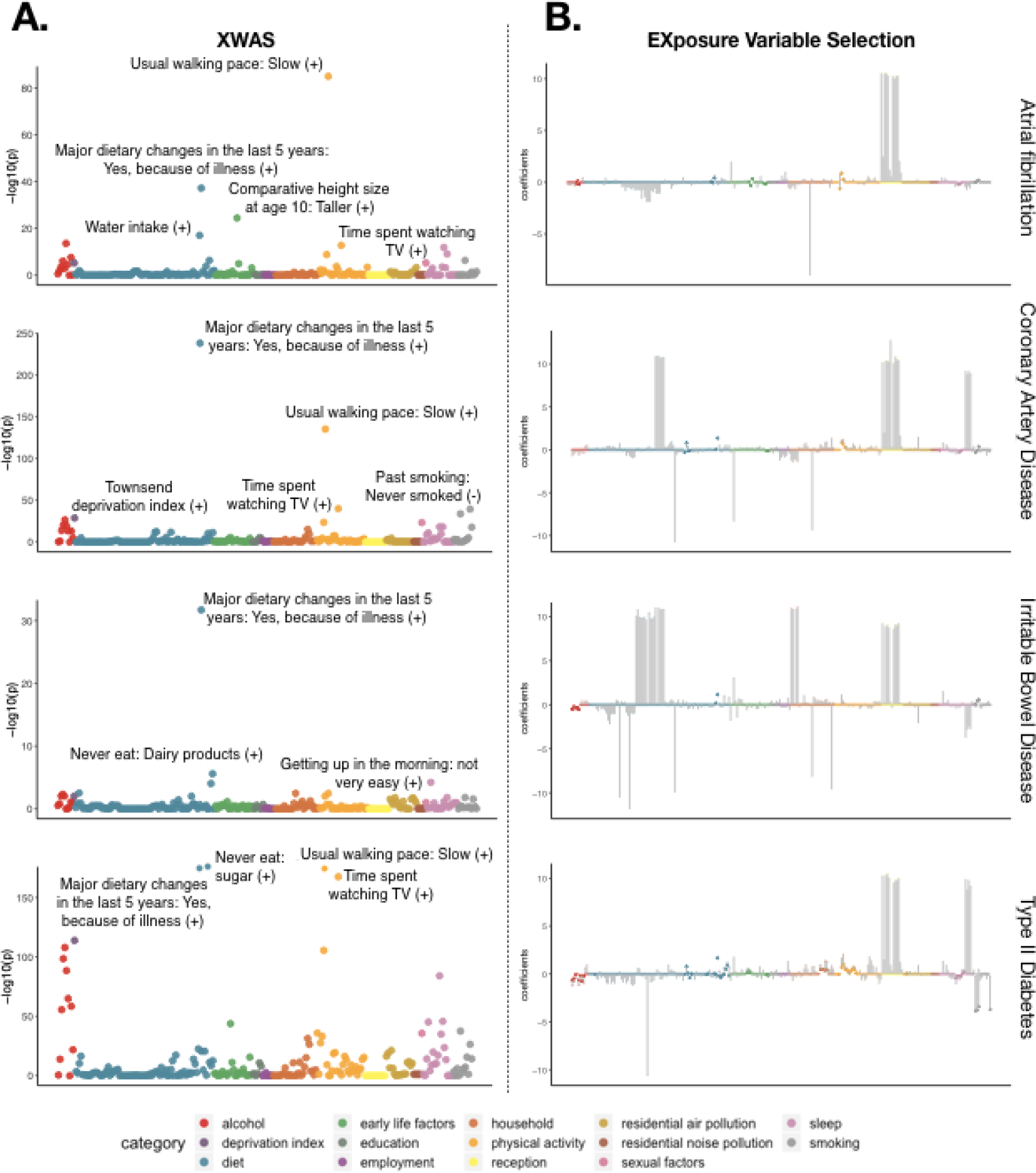
EXposure Wide Association Study (XWAS) and variable selection of each disease. A) eXposure-wide Association Study findings. Each exposure variable was regressed to each disease in group A with sex, age, and PC1-40 as covariates. The y-axis is the negative log of the *p*-value after FDR adjustment. The top most significant responses are labeled, with the direction of correlation in reference to the baseline response (Supplementary Table 1) in parenthesis. B) For each disease, we conducted variable selection on the significant exposure variables from XWAS in cohort B. The XWAS coefficients from cohort A are represented by a triangle, while the new weighted beta coefficients from the stepwise regression in cohort B are represented by a circle. Variables eliminated from selection are more transparent compared to those that were not. EXposure variables are colored by category and are ordered in the same way in both A and B.

We executed the variable selection step in group B (Figure 1). Depending on the disease, validation sample sizes ranged from 32,862 (T2D) to 76,966 (IBD) (Table 1). After selection, the number of significant exposures independently associated with each disease dropped to eight for AF and CAD (8.33%), three for IBD (3.12%), and 14 for T2D (14.58%) (Methods, Figure 2B, Supplementary Tables 2-5). The direction of effect was mostly consistent in the responses that were retained (78.95%, 84.21%, 81.82%, and 86% of responses for AF, CAD, IBD, and T2D, respectively). Variables retained for the diseases included indicators such as alcohol consumption, diet, early life factors (e.g. comparative body size at age 10), household income (e.g., average total household income before tax), physical activity, air pollution, sleep, and smoking activity. In the final logistic regression model with the selected variables, at least one response from each category was associated (*p*<0.05) with the disease. In our testing set (group C), the PXS, which encompasses non-genetic factors/exposures independently associated with disease, had a higher performance (defined as the maximum area under the receiver-operator curve [AUC]), than that of any single exposure variable (Figure 3). Compared to the average AUC of individual exposures, the AUC of PXS for AF, CAD, IBD, and T2D increased by 4.6%, 20.7%, 13.2%, and 22.0%, respectively (Figure 3).

**Table 1:** Disease prevalence. The number of individuals with complete exposure responses in validation and testing sets for each of the four diseases. (AF: atrial fibrillation, CAD: coronary artery disease, IBD: inflammatory bowel disease, T2D: type 2 diabetes.)

|  | Validation (Group B) |  | Testing (Group C) |  |
| --- | --- | --- | --- | --- |
|  | Cases/Total | Prevalence | Cases/Total | Prevalence |
| **AF** | 1,898/47,918 | 3.96% | 9,257/193,609 | 4.78% |
| **CAD** | 1,915/47,142 | 4.06% | 9,612/188,456 | 5.1% |
| **IBD** | 715/76,251 | 0.94% | 2,176/228,345 | 0.95% |
| **T2D** | 1,542/31,320 | 4.92% | 5,544/97,862 | 5.67% |

**Figure 3:**
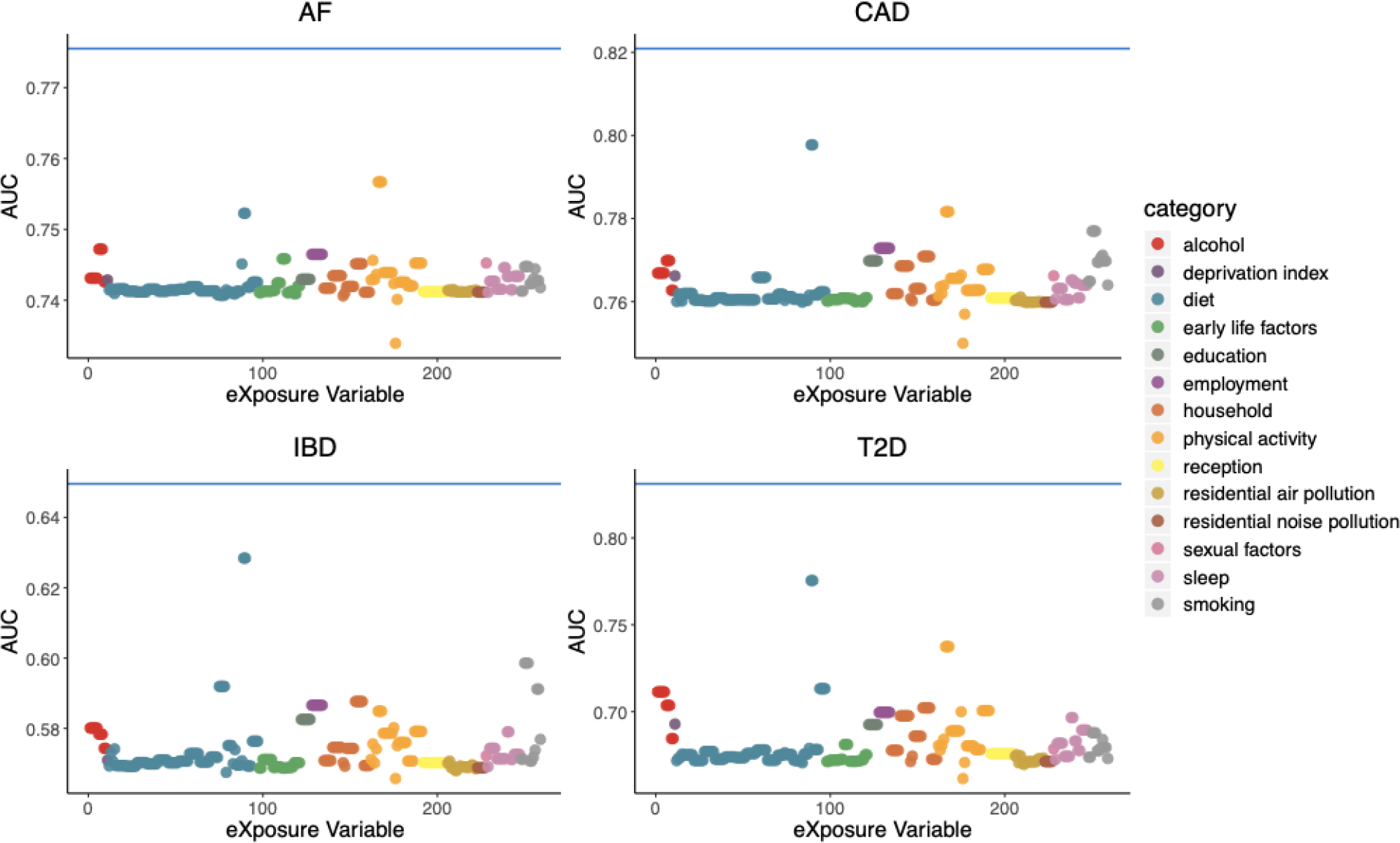
Combined effect of exposures is greater than any exposure alone. The blue horizontal line represents the AUC of PXS in the testing set with PXS, sex, and age as independent variables in group C. Each point is the AUC of an individual eXposure adjusted by age and sex. The eXposure variables are colored by category and are ordered in the same way in Figure 2.

### Additive non-genetic/eXposure factors play a more important role than genetics in disease prediction

We compared the matched case-control area under the ROC curve (AUC) for PDS, PXS plus PDS, PRS plus PDS, and combined scores (PDS+PRS+PXS) in cohort C (Table 1, Figure 4, Supplementary Table 7). In all diseases except for IBD, PXS prformed better than PRS. T2D had the largest discrepancy in performance (PDS AUC of 0.673 [0.663-0.683], PRS AUC of 0.711 [0.702-0.720], PXS AUC of 0.828 [0.821-0.836]). A “combined” score of the sum of PDS, PXS, and PRS performed better than any score alone (AUC of 0.834 [0.827-0.841] in T2D).

**Figure 4:**
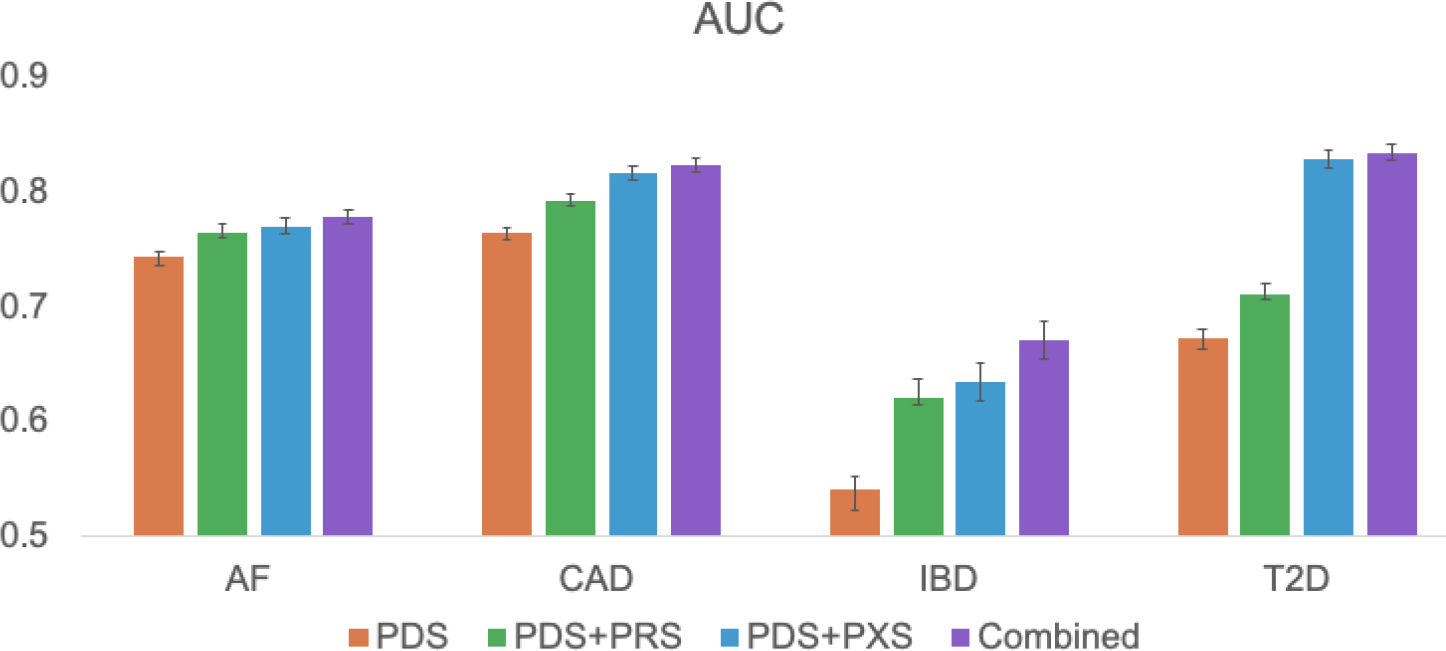
AUC of each score. Error bars represent the 95% confidence interval for each disease model. (PDS: poly-demographic score, PRS: polygenic risk score, PXS: poly-exposure score, AF: atrial fibrillation, CAD: coronary artery disease, IBD: inflammatory bowel disease, T2D: type 2 diabetes.)

Next, we compared the ability of exceptional tails of each score to identify individuals at substantially greater odds of disease (as in Khera et al.^3^). We found that the top one percent of PXS had much higher prevalence and odds of disease than PDS or PRS in all four diseases (Supplementary Figure 1, Table 2). In all diseases except for IBD, the median of the PDS was greater than PXS in the cases compared to controls (Supplementary Figure 2). Taking T2D as an example, we found that individuals in the top one percent of PXS had a greater than 15 fold greater odds of T2D relative to the remaining population, whereas individuals in the top one percent of PRS had a 3 fold greater odds relative to the remaining population (Table 2). The top percentile of PXS also had a much higher prevalence of T2D (53.27%) compared to PRS (15.24%) (Figure 5). We also found that the median percentile of PDS and PXS was greater than PRS in T2D cases versus controls (73%, 89%, and 67%, respectively) (Figure 5).

**Table 2:**
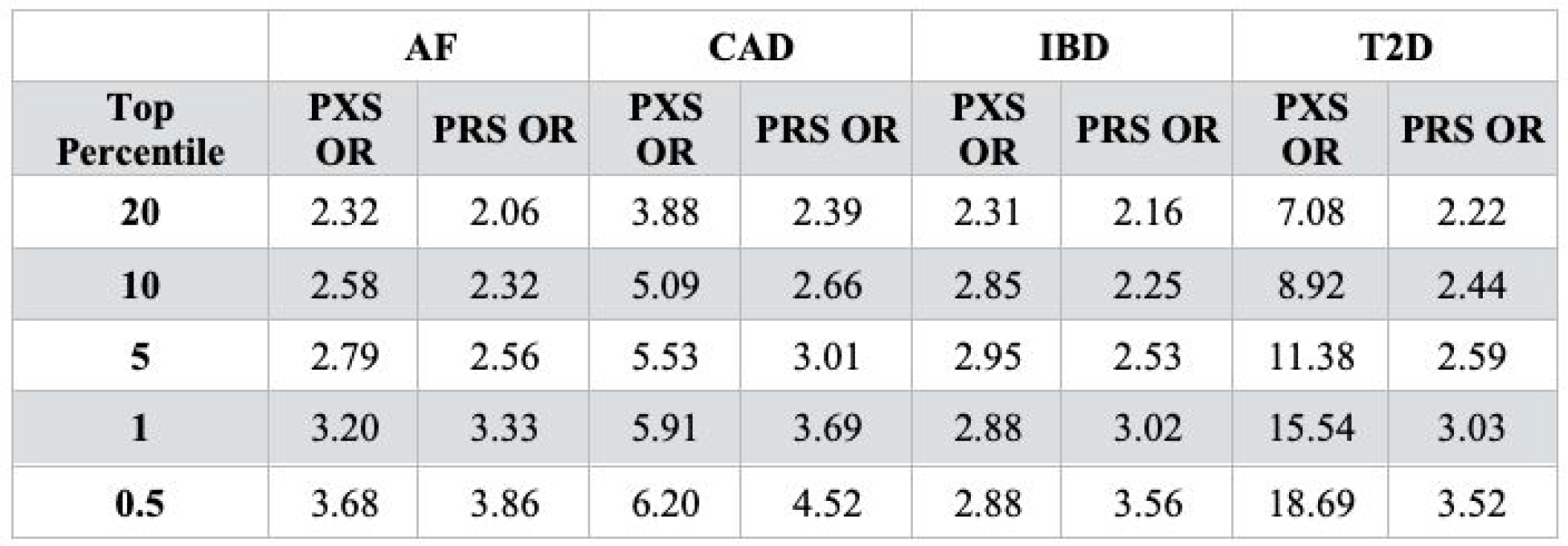
Odds ratios (ORs) of disease for the top percentile of the PRS, PXS, and PDS. ORs of the top percentiles compared to the remaining population in group C for each disease (e.g., the last row represents OR for the top 0.5% of the population versus the remaining 99.5% of the population). (AF: atrial fibrillation, CAD: coronary artery disease, IBD: inflammatory bowel disease, T2D: type 2 diabetes.)

|  | AF |  | CAD |  | IBD |  | T2D |  |
| --- | --- | --- | --- | --- | --- | --- | --- | --- |
| Top Percentile | PXS OR | PRS OR | PXS OR | PRS OR | PXS OR | PRS OR | PXS OR | PRS OR |
| 20 | 2.32 | 2.06 | 3.88 | 2.39 | 2.31 | 2.16 | 7.08 | 2.22 |
| 10 | 2.58 | 2.32 | 5.09 | 2.66 | 2.85 | 2.25 | 8.92 | 2.44 |
| 5 | 2.79 | 2.56 | 5.53 | 3.01 | 2.95 | 2.53 | 11.38 | 2.59 |
| 1 | 3.20 | 3.33 | 5.91 | 3.69 | 2.88 | 3.02 | 15.54 | 3.03 |
| 0.5 | 3.68 | 3.86 | 6.20 | 4.52 | 2.88 | 3.56 | 18.69 | 3.52 |

**Figure 5:**
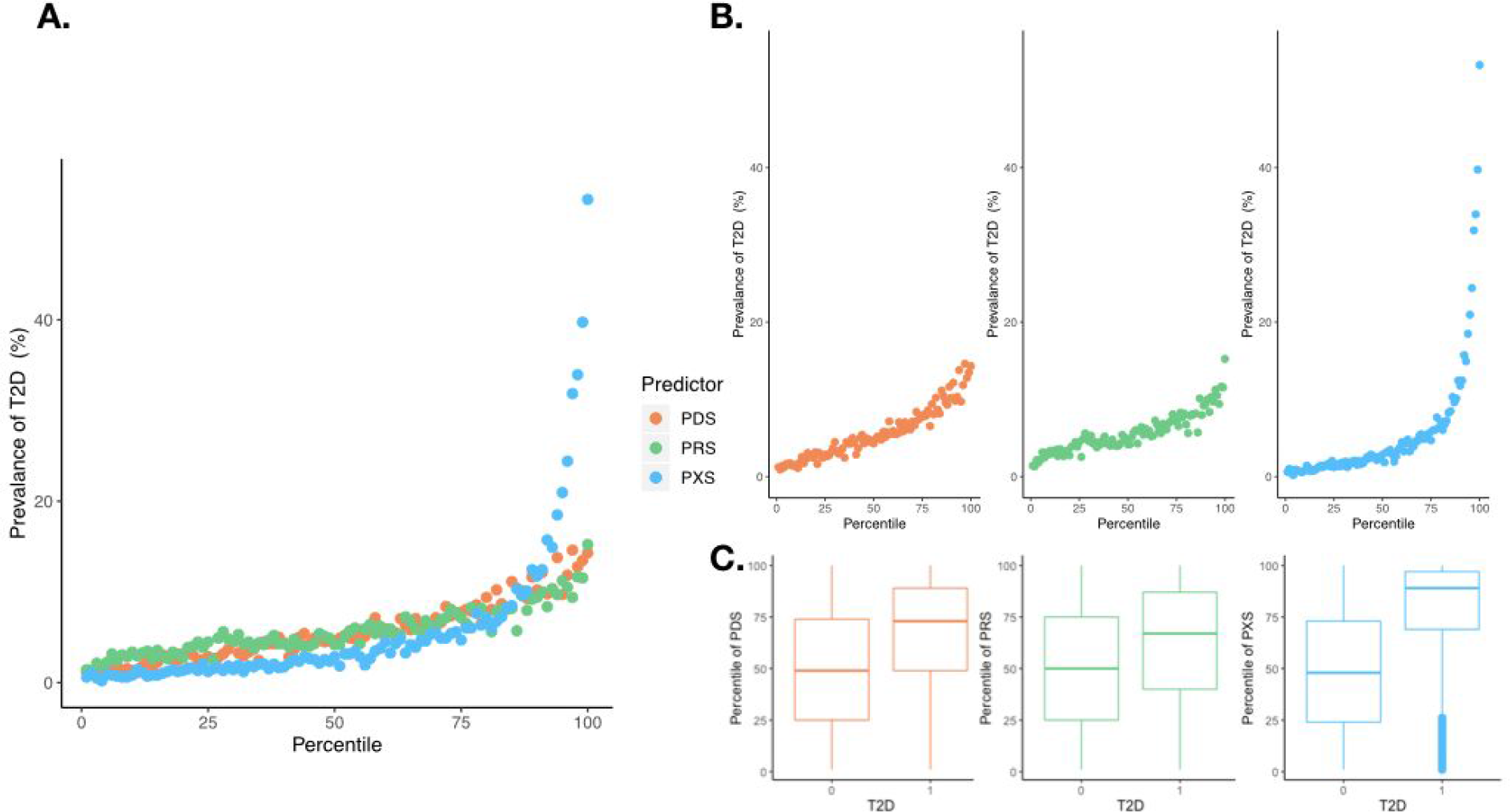
Relationship between PDS, PRS, PXS and Type 2 Diabetes. A) We binned participants in group C by their PDS (orange), PRS (green), or PXS (blue) percentiles, and we estimated the prevalence of T2D within each bin. B) Prevalence of disease versus PDS, PRS, and PXS. C) Distribution of PDS, PRS, and PXS in T2D cases and controls. For each boxplot, the middle horizontal line represents the median, and the top and bottom of each box represent the 25th and 75th percentiles. Dots represents outliers. The median PDS, PRS, and PXS percentile scores were 73, 67, and 89, respectively, while the median PDS, PRS, and PXS percentile scores for individuals without T2D were 49, 50, and 48, respectively. (PDS: poly-demographic score, PRS: polygenic risk score, PXS: poly-exposure score, T2D: type 2 diabetes.)

### PXS identifies individuals with high disease odds ratio but low PRS

Next, we measured the correlation between PXS and PRS in each disease to estimate the amount of independent information provided by each and aggregate “gene-exposure correlation”. The correlations between PXS and PRS were modest (Pearson correlation coefficient less than 0.1), but significantly non-zero, for each of the diseases except for IBD (Table 3). We investigated whether individuals with a high PXS would also have a high PRS. For each percentile of PXS, we calculated the average PRS percentile for individuals in that bin, and vice versa. Taking type 2 diabetes as an example, we found that the average PXS percentile of individuals in the top PXS percentiles were higher than that of the lowest percentiles (Figure 6). However, the average PRS percentile in the top percentiles of PXS were modest (an average PRS percentile of 59.27 in top 1% of PXS). Individuals in the top one percentile of PXS have a T2D odds ratio of 15.54 compared to the rest of the population (Table 2), but their risk for disease is masked if only PRS is considered. The general results were upheld in AF and CAD as well (Supplementary Figure 3).

**Table 3:** Correlation between PRS and PXS. (PCC: Pearson correlation coefficient, AF: atrial fibrillation, CAD: coronary artery disease, IBD: inflammatory bowel disease, T2D: type 2 diabetes.)

|  | *p*-value | PCC |
| --- | --- | --- |
| **AF** | 8.88E-32 | 0.027 |
| **CAD** | 1.99E-111 | 0.052 |
| **IBD** | 0.131 | 0.003 |
| **T2D** | 1.90E-110 | 0.071 |

**Figure 6:**
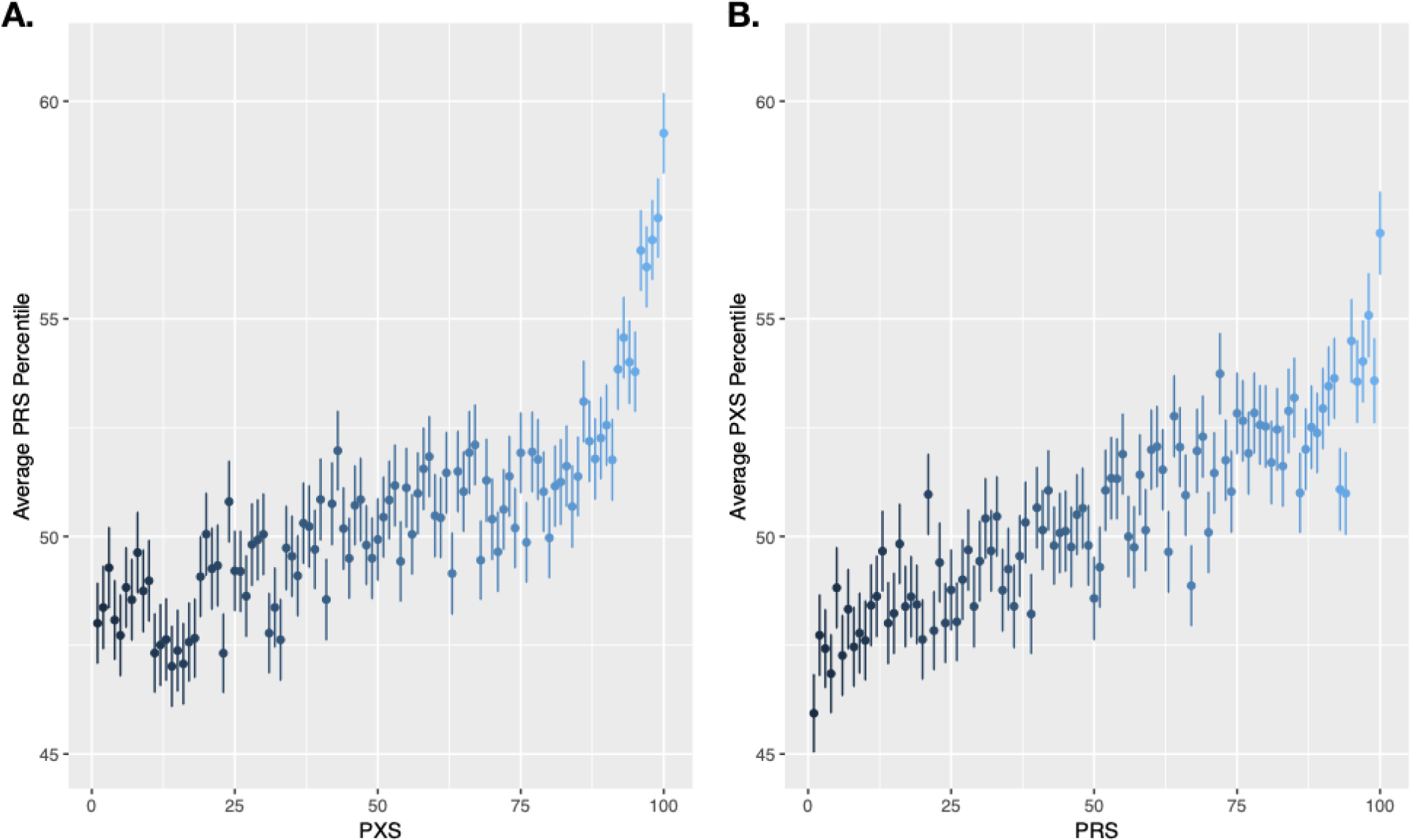
PXS identifies individuals with high odds for individuals with genetically low risk for T2D. A) The average PRS percentile of individuals in each PXS percentile. B) The average PXS percentile of individuals in each PRS percentile. Error bars represent the standard error of the data. (T2D: type 2 diabetes, PXS: poly-exposure score, PRS: polygenic risk score)

## DISCUSSION

Common diseases are well recognized to be due to a combination of genetic and environmental exposure factors. However, while thousands of genetic loci have been identified to be associated with complex diseases, nothing comparable has been executed for environmental exposures. Furthermore, researchers combine thousands of genetic markers through a polygenic risk score (PRS) using methods that accounts for linkage between SNPs^10^, but studies on exposures remain, for the most part, focused on a small set of exposures at a time without much consideration for dense correlation between exposures.

In our study, we term the poly-exposure score (PXS), which encompasses information from many exposures simultaneously. Analogous to PRS, which are estimated from GWAS summary statistics^10^, PXSs can be derived from multivariate XWAS summary statistics. PXSs are simple to estimate and translate for clinical care, and much of the non-genetic exposure information is less invasive to collect than genetic information. To demonstrate the simplicity of the PXS score, we have created an interactive resource (http://apps.chiragjpgroup.org/pxs/) to calculate PXS for each of the diseases.

Further still, our investigation serves as a reminder of the premium importance of demographic factors, such as age and sex, in risk prediction. While most risk models or associative approaches include demographic variables, new models are often presented without a comparison of simpler “baseline” models (e.g. Khera et al.^3^), which may mislead their consumers and pose a challenge for replication.

The results of our study have many implications. First, we show that PXSs are able to predict disease outcome better than any single exposure alone (Figure 3), but PDS, or age and sex, describe most of disease risk. Taken together, this supports the paradigm that the exposome, which refers to the totality of environmental (non-genetic) exposures, play a large role in diseases^11,12^. Rather than studying the relationship between individual exposure variables with a disease, we should investigate the effects of “poly-exposures” through PXSs.

We also demonstrate that PXS performs as well as, if not better than, PRS in identifying individuals with the highest odds for diseases. For example, in type 2 diabetes, individuals in the top one percent of PXS had an odds ratio of 15.5 (versus the rest of the population), whereas individuals in the top one percent of PRS had an odds ratio of 3.03 (Table 2). Furthermore, we found that type 2 diabetes occurs with a staggering prevalence of 53.27% in the top percentile of PXS, compared to prevalences of 15.24% and 14.31% in the top percentile of PRS and PDS, respectively (Figure 5). In T2D, the median PDS, PRS, and PXS percentile scores were 73, 67, and 89, respectively, while the median PDS, PRS, and PXS percentile scores for individuals without T2D were 49, 50, and 48, respectively, in individuals without T2D.

Most importantly, our results suggest that the PXS can capture crucial information missed by PRS. While PXS and PRS are significantly correlated with each other, individuals with a high PXS may not have a high PRS, and vice versa (Figure 6). PXS is able to identify individuals with high disease odds ratio but low PRS. Taken together, our results indicate that while PRS is useful for screening individuals with high genetic risk, for most individuals PRS conveys only a fraction of the story in total disease risk.

Through our XWAS analysis, we were also able to replicate several documented epidemiological associations between diseases and exposures. In type 2 diabetes (T2D), observational studies have implicated “lifestyle” exposures, which include behaviors such as physical inactivity^13^ and smoking^14^, to be associated with T2D. Further, environmental exposures such as air pollutants^15^ have also been positively associated with T2D; however, their associations are weaker (smaller effect sizes) than smoking or physical activity. In our analysis, the relationship between T2D and higher air pollution (PM10) and physical inactivity (“Time spent watching TV”, “Frequency of stair climbing in the past 4 weeks”, etc.) were confirmed. We also confirmed previous results that showed positive associations of smoking with AF^16^ (“Past smoking history”), T2D^14^ (“Ever smoked”), and CAD^17^ (“Past tobacco smoking”). The relationship between smoking and IBD is complex^18^, but our results suggest a positive association between IBD and history of smoking (“Past smoking history”). We also uncovered, to the best of our knowledge, novel associations between exposures and each of the diseases. Notably, the response “No” for the variable “Maternal smoking around birth” was significantly and negatively associated with AF, CAD, and T2D in reference to the response “Yes” (Supplementary Table 1).

Using time-dependent and self-reported non-genetic factors and exposures to predict diseases has several limitations. In many cases, multiple exposure variables in the UK Biobank are correlated and/or measure similar responses. For example, there are several variables in the UK Biobank that measure alcohol consumption, including “Alcohol drinker status”, “Alcohol intake versus 10 years previously” and “Alcohol intake versus 10 years previously”. Theoretically, the forward stepwise regression should eliminate variables that contain correlated or redundant information. Only variables that are independently associated with the disease are retained. In the case of T2D alone, both “Ever smoked” and “Past smoking history” were both retained, but the responses coded non-redundant information, i.e. “Ever smoked: No” and “Past smoking history: Never smoked” were reference groups.

Relatedly, the most significant challenge in observational exposure studies is the deduction of direction of causality or potential confounding variables. For example, it is possible that the significant associations of the response “Major dietary changes in the last 5 years: Yes, because of illness” with all four diseases is explained by the onset of each disease or possibly another illness (comorbidity). Further, it is hypothesized that confounding and model misspecification occurs at higher rate in non-genetic vs. genetic studies^19^. Third, while easy to measure, some of the exposures considered included self-reported variables, such as diet, which may be prone to measurement error and recall bias^20^. If these errors occur at random across all variables considered in the PXS, the association sizes and PXSs will be diluted. If, on the other hand, individuals in the cases versus controls report their intakes differently, the PXSs will also be directionally biased. It is less clear how PXS will be affected if the types of the errors are different (both random and differential with respect to the exposure or disease) across the variable inputs. Further analysis using methods like Mendelian randomization may address confounding and reverse causality at biobank scale^21^ for individual indicators, but these issues will need to be evaluated for groups of exposure factors as well.

We only considered exposure variables if they contained less than 10% missing data. Increasing data completeness of variables, or imputing exposure information, would be valuable to eventually employ machine learning techniques for modeling. We included the responses “Do not know” and “None of the above” for categorical responses in our exposure study, as we believed these responses may code additional information and that the removal of individuals who did not know an answer would bias our findings. However, we found that many of these responses were not significant in either XWAS or post-stepwise variable selection. Still, we included all responses of an exposure variable for PXS derivation if at least one of the responses of the variable was significant in order to provide an accurate reference group for each categorical responses.

An inherent challenge to environment/non-genetic and genetic studies is that they are often examined in isolation. For example, genetic and exposure factors may be correlated, a phenomenon known as “gene-environment correlation”. To this end, we found that PXS and PRS had a modest but significant correlation with each other in all diseases except IBD (Table 2). It is hypothesized that the gene-environment interaction plays a large role in complex diseases^22^, but its effect on phenotypic variation is widely debated^23,24^. Further analysis is needed to elucidate the relationship between environment, genetics, and their interactions^22,25,26^.

Because the UKB consists of primarily individuals with European ancestry, we limited our analysis to only white participants. It is difficult to extrapolate these results to other ethnic populations; Martin et al. showed that polygenic risk scores derived from Eurpoean GWASs were biased when applied to more diverse populations^27^. Furthermore, exposure disparities, such as socioeconomic status^28^, education attainment^28^, pollution^29^, and smoking^30^, are correlated with ethnicity. Therefore, there is a clear need for more diverse populations in both genetic^31^ and environmental exposure studies. A few notable studies exist or will be available in the future, such as the Malaysian Cohort Study^32^, All of Us Project^33^ and Kadoorie Biobank^34^. To capture the comprehensive variation of environment and genetics in diseases -- and to test the utility of precision medicine -- investigations in other populations will be instrumental.

## METHODS

### Study Governance

The UK Biobank (UKB) is a biobank of UK participants to examine the role of genetics and environmental exposures in human health. The UKB resource comprises of 502,655 participates between 40-69 years of age at the time of recruitment between 2006 and 2010. Participants attended one of 22 assessment centers across England, Scotland, and Wales, where they completed touchscreen and nurse-led questionnaires, had physical measurements taken, and provided biological samples. The study collected extensive data from questionnaires, interviews, health records, physical measures, biological samples, and imaging. UKB also collected information on individual background and lifestyle, cognitive and physical assessments, sociodemographic factors and medical history. UK Biobank has ethical approval from the NHS National Research Ethics Service. All participants provided informed consent.

### Quality Control of Data

We divided the white participants with complete demographic data (sex, age, PCs 1-40) were into groups A, B and C (N=459,613) (Figure 1, Table 1). These included individuals who self reported ethnicities of “British”, “Irish” and “Any other white background”. Group A (training) had 104,624 individuals, group B (validation) had 104,588 individuals, and group C (testing) had 250,401 individuals to begin with. For each of the four diseases, we conducted the initial univariate XWAS analysis and calculation of poly-demographic score coefficients in cohort A. We conducted exposure variable selection and coefficient adjustment of poly-exposure score with stepwise regression in cohort B and validated all three scores in cohort C.

We classified exposure variables as indicators of physiological state, environmental exposure and self-reported behavior. These were variables in the categories ‘Reception’, ‘Employment’, ‘Sociodemographics’, ‘Lifestyle and environment’, ‘Estimated nutrients yesterday’, ‘Early life factors’, ‘Typical diet yesterday’, ‘Meal type yesterday’, ‘Spreads/sauces/cooking oils yesterday’, ‘Alcoholic beverages yesterday’, ‘Hot/cold beverages yesterday’, ‘Cereal yesterday’, ‘Milk/eggs/cheese yesterday’, ‘Bread/pasta/rice yesterday’, ‘Soup/snacks/pastries yesterday’, ‘Meat/fish yesterday’, ‘Milk/eggs/cheese yesterday’, ‘Vegetarian alternatives yesterday’, ‘Fruit/vegetables yesterday’, ‘Residential air pollution’, ‘Residential noise pollution’. There were 206 unique variables in total. From these, we considered only the variables that had data for >90% of the participants as potential correlates (referred to as ‘factors’). There were 96 variables that remained.

We considered categorical responses of “Prefer not to answer” and continuous responses of −10,−3, and −1 missing data, which were removed from regression analysis at each stage of the exposure analysis. The number of individuals that remained for validation and testing can be found in Table 1.

### Phenotype ascertainment

UKB contains self reported data during an interview with a trained nurse as well as International Classification of Diseases (ICD-9 and ICD-10) diagnostic codes and Office of Population Censuses and Surveys (OPCS-4) surgery codes recorded across all episodes of hospital visit. Many codes represent the same overarching disease. For example, ICD-9 code 4274 codes of atrial fibrillation flutter, while ICD-10 I48.1 code for persistent atrial fibrillation and I48.2 code of chronic atrial fibrillation. Therefore, we based our grouping system to that found in Khera et al.^3^ of combined multiple self reported data, ICD9/10, OPCS-4 codes for each of the ascertainment of each disease.

Atrial fibrillation (AF) ascertainment was based on either self reported atrial fibrillation, ICD-9 427.3, ICD-10 I48.X, or OPCS-4 K57.1, K62.1, K62.2, K62.3, K62.4. Coronary artery disease (CAD) ascertainment was based on either self reported heart attack/myocardial infarction, ICD-9 410.9, 411.9, or 412.9, ICD-10 I21.X, I23.1, I23.2, I23.3, I23.6, I23.8, I24.1, I25.2, or OPCS-4 K40.X, K41.X, K45.X, K49.X, K50.2, or K75.X. Inflammable bowel disease (IBD) ascertainment was based on either self reported inflammatory bowel disease, ICD-9 555.X or ICD-10 K51.X. Type 2 diabetes (T2D) ascertainment was based on either self reported type 2 diabetes or ICD-10 E11.X.

In total, there were 22,846 individuals with atrial fibrillation, 25,909 individuals with coronary artery disease (CAD), 4,575 individuals with inflammatory bowel disease (IBD), and 30,108 individuals with type 2 diabetes (T2D).

### Poly-Demographic Score

We considered sex, age, and first 40 genetic principal components (PCs) to be demographic variables. Genetic PCs were included since they provide information on geographical location and ancestral background^35^. For each of the four diseases, we calculated the coefficient of each demographic factor against the phenotype indicator in group A in 42 separate logistic regressions.

We used the coefficients from logistic regression as an estimate of the direction of effect of the dependent variables. For example, a negative coefficient would indicate a negative correlation between the variable and the disease.

We calculated the PDS of individuals in group C in the following way:

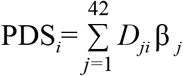

Where PDS of the individual i is equal to the weighted sum of the individual’s 42 demographic information. *D*_*ji*_ is the incident of demographic variable *j* for individual *i*. and β_*j*_ is the coefficient of variable *j* from the logistic regression.

### Executing a X-Wide Association Study (XWAS) and estimation of the poly-exposure score

Analogous to PRS, PXS can be calculated using summary statistics from exposure wide association study (XWAS)^8^. For each of the four diseases separately, we first conducted XWAS in group A. We associated each of 96 non-genetically measured environmental exposure, physiological state, and self-reported behavioral factors with the disease while adjusting for age, sex and first 40 principal components. Specifically, we modelled each of the factors as an independent variable, and disease (case/control) as the dependent variable while adjusting for covariates in 96 separate logistic regression models. To maximize sample size, individuals with missing data for each factor was removed for prior to running its regression model. We used the Benjamini-Hochberg False Discovery Rate (FDR)^36^ and deemed an FDR adjusted *p*-value of < 0.05 as significant. Variables that had at least one significant response were retained.

To minimize statistical interaction of the exposure variables, we used a modified forward forward stepwise regression method to select for independent variables. Forward selection is a method in which variables are iteratively added into a multivariate model based on a criterion to find the subset of variables in the dataset that results in the best performing model (as ascertained by R^2^). We conducted forward selection on the variables from the XWAS that passed FDR adjusted *p*-value <0.05 in group B samples with individuals with missing data removed. More specifically, we begin with a logistic model of the disease with the most significantly associated exposure variable and age, sex and PC 1:40 as covariates. We then progressively added variables that pass the threshold to the multivariate model. The coefficients of categorical variables represent the difference in effect of each class to the reference class. These coefficients were retained for the PXS calculation.

PXS was calculated in group C in a similar fashion as PDS:

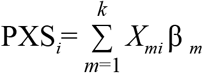

Where PXS of individual *i* is the weighted sum of the individual’s exposures. β_*m*_ is the coefficients from the stepwise multivariate logistic regression model. *X*_*mi*_ is the incident of exposure variable *m* for individual *i*. Individuals with missing data in any of the remaining exposure variables were removed from group C.

### Calculating the polygenic risk score for UK Biobank participants using weights from Khera et al

We calculated polygenic risk scores (PRS) for UKB participants with weights published by Khera et al^3^. In short, Khera et al. estimated weights summary statistics from recent GWAS studies in participants of European ancestry. They generated candidate scores for a subset of the UK Biobank participants using the LDPred algorithm^10^, and selected the best score based on maximum AUC in a logistic regression model with disease as the outcome and the score, age, sex, PC1:40 as covariates.

With the aforementioned weights, we calculated the PRS of individuals in group C using the built in allelic scoring procedure of PLINK (--score)^37^. PLINK takes the sum of the number of each reference allele multiplied by the weighted coefficient of the allele across all alleles.

### Estimating disease odds ratio (OR) and area under the curve (AUC) in the UK Biobank testing dataset

For each disease, we calculated the area under the curve (AUC) in a logistic regression model with either PDS, PDS+PXS, PDS+PRS, or PDS+PXS+PRS. Since demographic variables are typically considered as covariates in genetic and environmental exposure studies, we included PDS in PRS versus PXS comparison. To account for the disease and control sample size imbalance, we averaged 100 AUC’s (bootstrapped 1000 iterations) calculated in sampled populations of group C where the number of disease controls matched cases. Odds ratios for each score were derived by comparing the top 20%, 10%, 5%, 1% and 0.05% of the distribution with the remaining individuals in a logistic regression with sex, age, and PC1:40 as covariates.

We placed individuals into 100 bins by their PDS, PXS or PRS percentiles for each disease. Within each bin, we then calculated the prevalence of disease cases. We estimated the average PRS and PXS percentiles within each PXS and PRS bin, respectively. We also estimated the Pearson correlation coefficient and *p*-value between PRS and PXS in each disease to measure gross PRS and PXS correlation.

We fit all logistic regression models with the ‘glm’ function in R. AUCs and the 95% confidence intervals were calculated using the ‘auc’ function in the pROC package^38^ of R. We used the “p.adjust” function of the base stats R package^39^ for Benjamini-Hochberg False Discovery Rate (FDR) adjustment for multiple tests.

### Web Resources

Code repository: https://github.com/yixuanh/poly-exposure-score

PXS calculator: http://apps.chiragjpgroup.org/pxs/

## ACKNOWLEDGEMENTS

Our analysis was conducted using the UK Biobank resource via application number 22881. We would like to thank all the volunteers who participated in this project. This work was supported by the Bioinformatics and Integrative Genomics training grant from the National Institutes of Health NHGRI under award number T32HG002295, the National Institutes of Health NIEHS under award numbers R00ES23504 and R21ES205052, NIAID under award numbers R01AI12725003, the UK Biobank Early-Career Researcher Award (to Y.H.), and the National Science Foundation under award number 1636870.

