## Supplementary Figures for "Poly-Exposure and Poly-Genomic Scores Implicate Prominent Roles of Non-Genetic and Demographic Factors in Four Common Diseases in the UK"

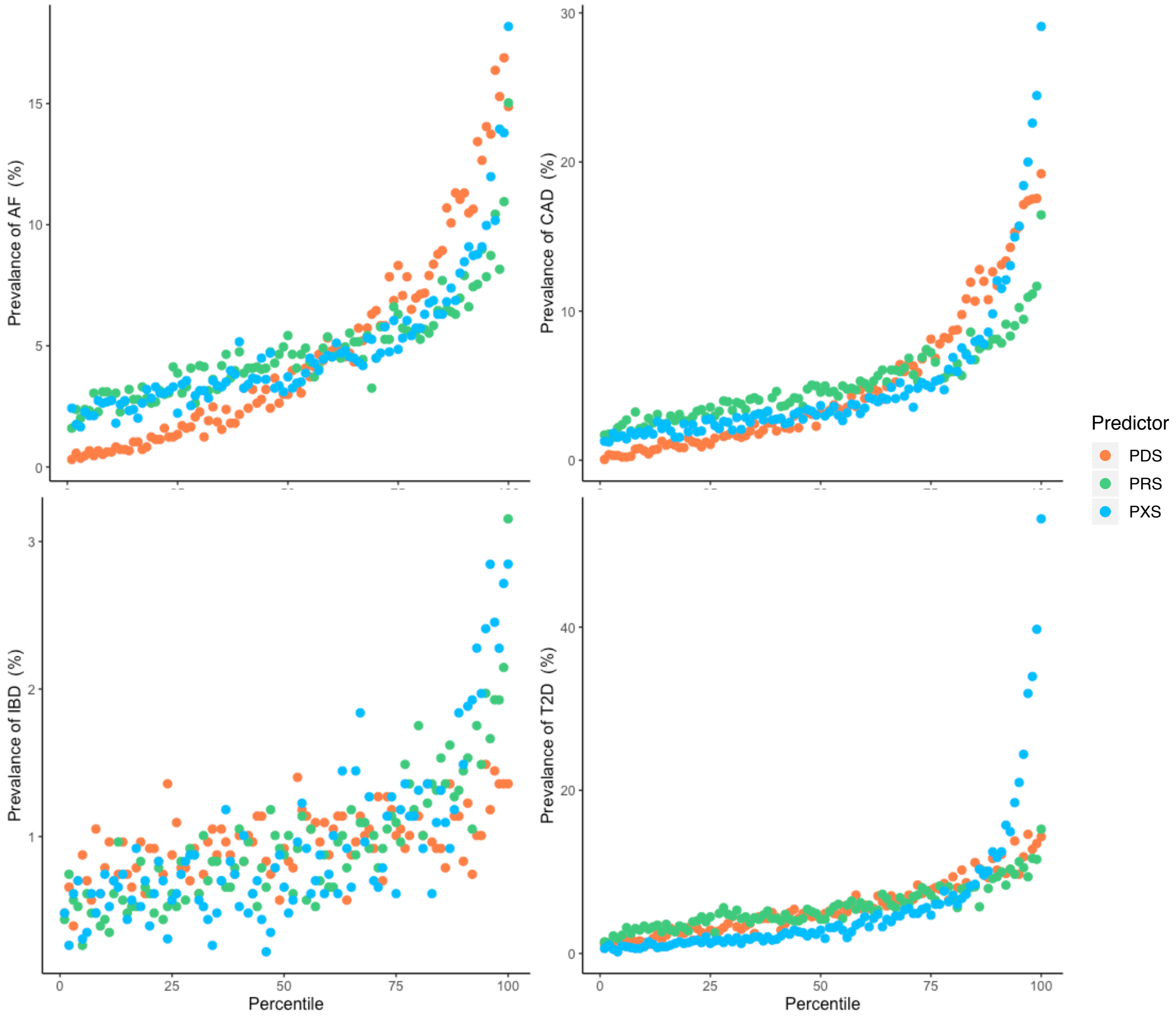


**Supplementary Figure 1: Relationship between PDS, PRS, PXS and diseases.** Individuals were binned into 100 groups based on their PDS (orange), PRS (green), or PXS (blue), and the prevalence of each disease within each group was calculated.

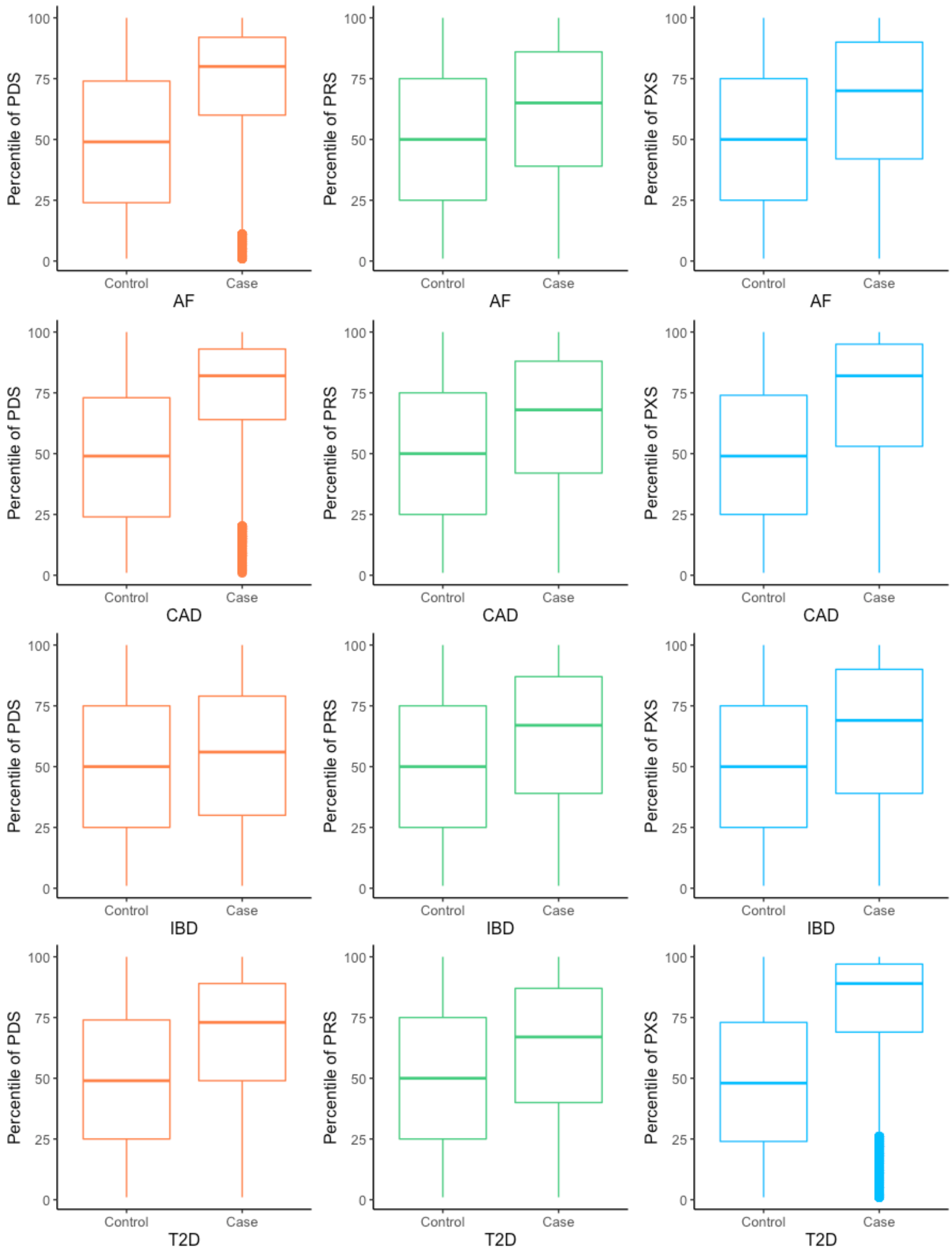
Percentile of PDS
Percentile of PRS
Percentile of PXS
Control
Case
AF
CAD
IBD
T2D
0
25
50
75
100

**Supplementary Figure 2. PDS, PRS, and PXS between disease cases and controls.** For each boxplot, the middle horizontal line represents the median, and the top and bottom of each box represent the 25th and 75th percentiles. Dots are represented as outliers.

AF

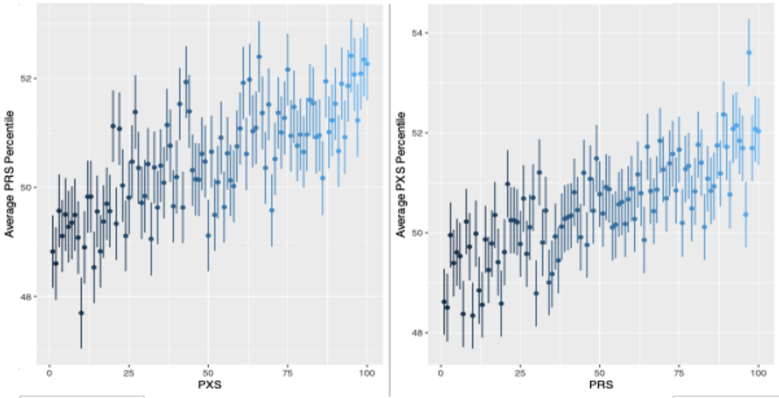


CAD

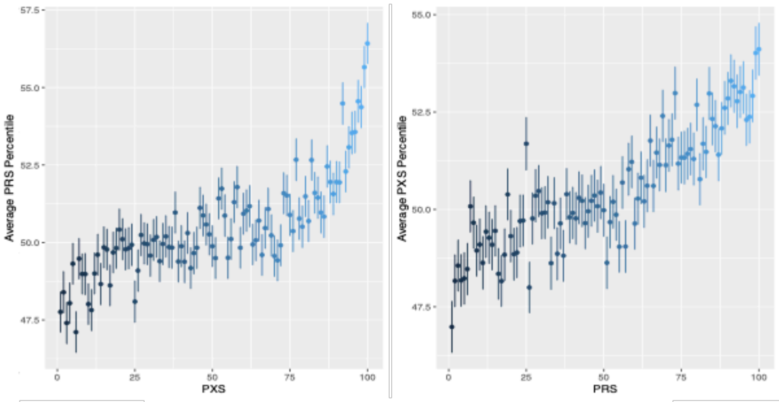


IBD

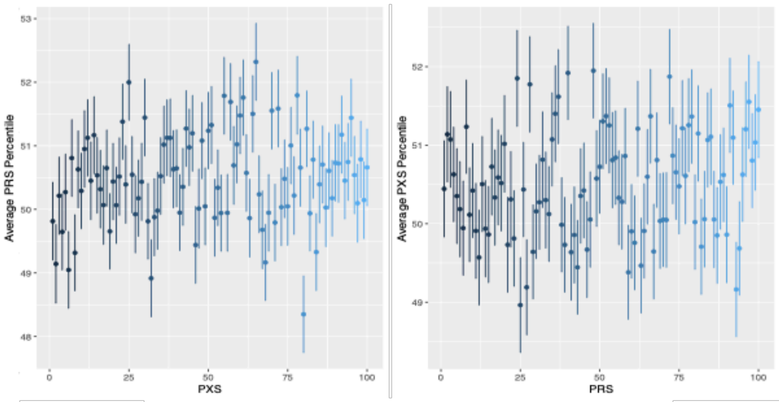


T2D

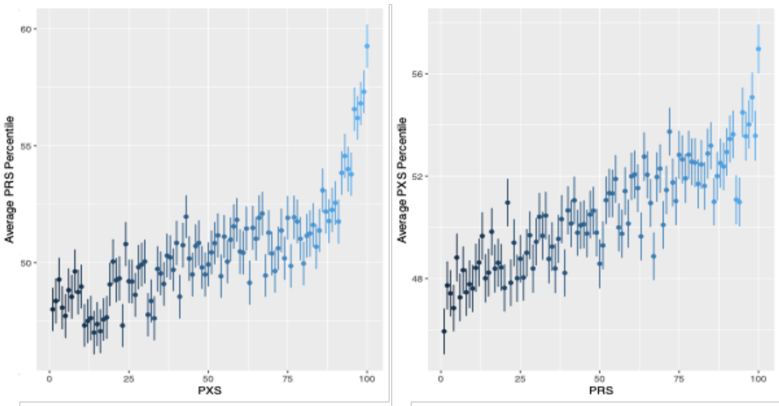

**Supplementary Figure 3. Correlation between PXS and PRS is diseases.** For each disease, the left plot shows the average PRS of individuals in each PXS percentile. The average PXS of individuals in each PRS percentile. Error bars represent the standard error of the data

**Supplementary Table 1. XWAS for AF, CAD, IBD, and CAD.** For each of the 96 exposure variables, we conducted an XWAS with each disease. We report the beta coefficient, AUC, p-value, and FDR-adjusted p-value for each association. AUC was determined in a logistic regression with sex, age, and PC 1:40 as covariates. Each entry is in references to the baseline response, indicated in brackets. The sample size for each XWAS is also shown.

**Supplementary Table 2. Weighted coefficients of exposure variables selected for PXS calculation in atrial fibrillation.** Using a modified step-wise regression, we selected for independent and significant variables associated with each disease. Eight variables were selected.

**Supplementary Table 3. Weighted coefficients of exposure variables selected for PXS calculation in coronary artery disease.** Using a modified step-wise regression, we selected for independent and significant variables associated with each disease. Eight variables were selected.

**Supplementary Table 4. Weighted coefficients of exposure variables selected for PXS calculation in inflammatory bowel disease.** Using a modified step-wise regression, we selected for independent and significant variables associated with each disease. Three variables were selected.

**Supplementary Table 5. Weighted coefficients of exposure variables selected for PXS calculation in type 2 diabetes.** Using a modified step-wise regression, we selected for independent and significant variables associated with each disease. 14 variables were selected.

**Supplementary Tables 6. Weighted coefficients of variables used for PDS calculation.** The coefficient and p-value of each variable to disease is shown.

**Supplementary Table 7: Comparison of PDS, PRS, and PDS.** The AUC was determined in in a logistic regression of each score with disease.
